# Heterologous expression in *E. coli* reveals the bicarbonate transporter BicA2 drives carbon uptake in marine *Prochlorococcus* spp

**DOI:** 10.64898/2026.05.23.727450

**Authors:** Loraine M. Rourke, Caitlin S. Byrt, G. Dean Price, Benedict M. Long

## Abstract

The widespread oceanic cyanobacterial *Prochlorococcus* genus is a major contributor to global carbon fixation, yet mechanisms enabling this lineage to elevate intracellular inorganic carbon as a substrate for photosynthesis remain unresolved. Cyanobacterial CO_2_ concentrating mechanisms typically rely on membrane-bound bicarbonate (HCO_3_ ^-^) transporters SbtA1, SbtA2, BicA and BCT1, and CO_2_-to-HCO_3_^-^ conversion uptake systems (CO_2_ pumps; NDH-I_3_ and NDH-I_4_), to elevate a cellular HCO_3_^-^ pool for use by Rubisco-containing carboxysomes. Evidence suggests *Prochlorococcus* harbours carboxysomes with a low-CO_2_-specificity Rubisco, implying a functional CCM dependent on active HCO_3_^-^ uptake. However, canonical CO_2_ pumps are absent, leaving distant HCO_3_ ^-^ transporter homologues, BicA2 and SbtA2, as prime candidates for HCO_3_^-^ transport in this group. Yet these have not been functionally characterised. Here we demonstrate that BicA2 from *P. marinus* CCMP1375 mediates Na^+^-dependent HCO_3_^-^ uptake in *E. coli*, while BicA2 from *P. marinus* CCMP1986 is inactive in its native form but acquired transport function through a single amino acid substitution during adaptive laboratory evolution. These findings confirm BicA2 as a low-affinity, Na^+^-dependent bicarbonate transporter with variable flux, revealing a previously uncharacterized CCM component in *Prochlorococcus*. This mechanistic insight reshapes our understanding of carbon acquisition strategies in the most abundant photosynthetic organism on Earth and highlights evolutionary plasticity in transporter function with implications for global biogeochemical cycles.

**Highlight:** This study provides the first mechanistic evidence of bicarbonate transport in *Prochlorococcus*, revealing functional and evolutionary flexibility in its CO_2_-concentrating mechanism components and reshaping our understanding of carbon acquisition in the most abundant marine photoautotroph.

## Introduction

*Prochlorococcus* spp., along with oceanic *Synechococcus* spp. of the α-cyanobacterial sub-group [1], are attributed with an estimated 25% of annual net CO_2_ fixation in marine environments [2]. *Prochlorococcus* are the smallest and most abundant single-celled photosynthetic organism [3,4]. They have some of the smallest genomes of all photosynthetic organisms [5], yet they are genetically diverse, with ∼50% of the pangenome conserved across species [6]. The remainder of the genome is quite diverse, most likely due to the variation in environmental niches in which they grow, including high-light (typically surface-water) and low-light (typically deep-water) adapted ecotypes [6].

Efficient inorganic carbon (Ci) accumulation to drive photosynthesis in cyanobacteria and other phytoplankton (for example diatoms and green micro-algae) is primarily attributed to their CO_2_-concentrating mechanisms (CCMs) that support high rates of photosynthetic CO_2_ fixation [7,8]. The CO_2_ concentration is increased in the immediate vicinity of the site of the CO_2_-fixing-enzyme, ribulose-1,5-bisphosphate carboxylase/oxygenase (Rubisco), which converts CO_2_ to sugars for net carbon gain [7,9,10]. As a key element of their CCMs, cyanobacteria have several CO_2_ and HCO_3_^-^transporters located on the thylakoid and plasma membranes (Figure 1), respectively, that actively accumulate bicarbonate (HCO_3_^-^) into their cells [11]. This elevated HCO_3_^-^pool then diffuses into protein-encapsulated micro-compartments called carboxysomes [7,12] where carboxysomal carbonic anhydrase (CA), co-located with Rubisco, inter-converts HCO_3_^-^ and CO_2_, increasing the available CO_2_ concentration for rapid carboxylation [13,14].

**Figure 1:**
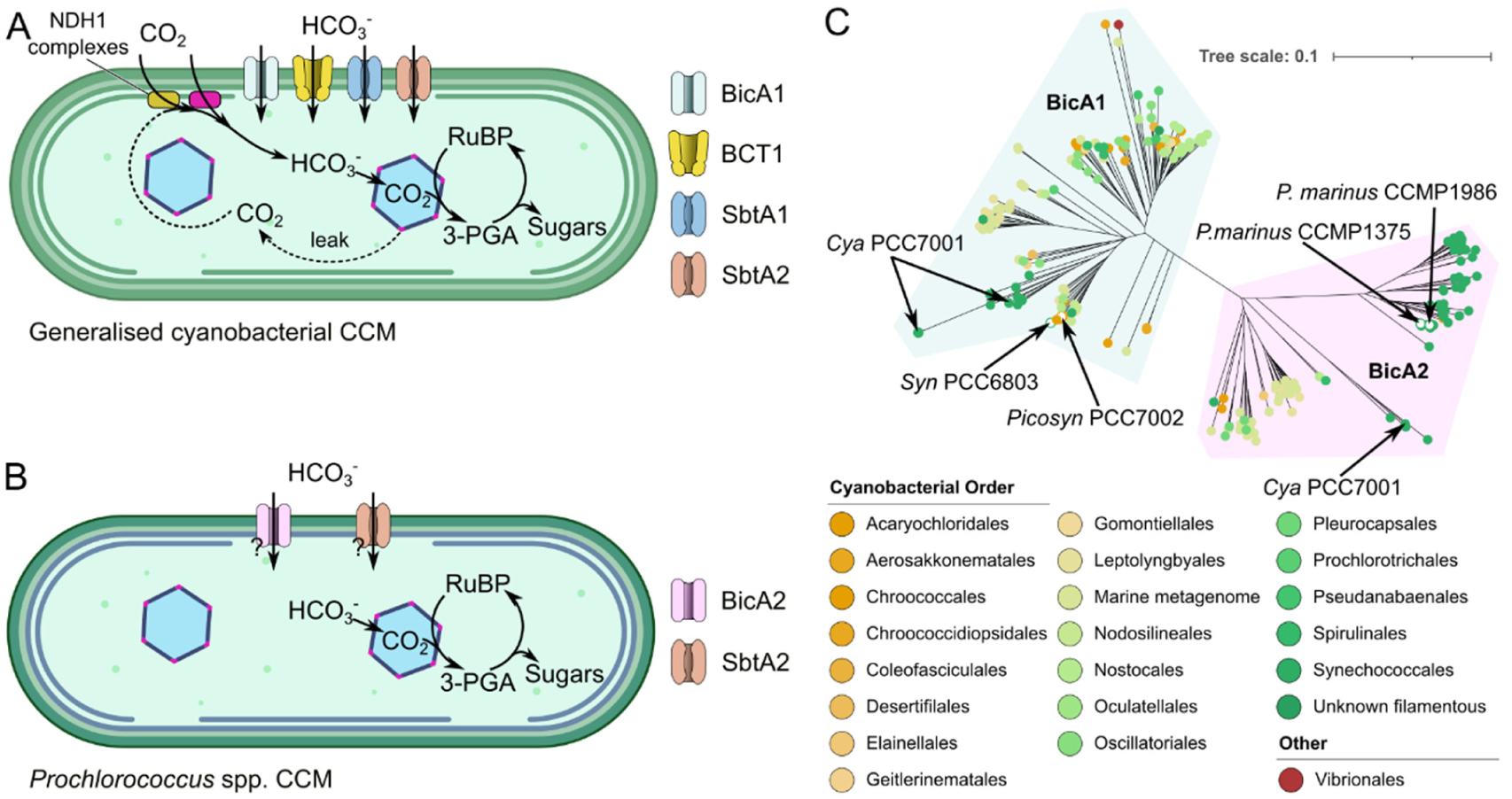
Inorganic carbon (Ci) transport in generalised cyanobacterial CO_2_ concentrating mechanism (CCMs) and *Prochlorococcus* spp. **(A)** CO_2_ concentrating mechanism components in most cyanobacteria is generalised to be driven by inorganic carbon (Ci) transport by up to four characterised bicarbonate transporters (BicA1, BCT1, SbtA1, SbtA2; [15–17,19,47,56]) and two CO_2_ to HCO_3_-conversion systems [NDH-1 complexes using ChpX or ChpY; 11,57] on the thylakoid membranes. These collectively supply a large cellular HCO_3_- pool that serves the Rubisco-containing carboxysome, where a specialised carbonic anhydrase enables elevation of CO_2_ concentrations for rapid CO_2_ fixation. **(B)** In *Prochlorococcus* spp., genes coding for members of the SbtA2 [19] and putative BicA2 transporters are proposed to enable efficient CCM function [29]. However, functional SbtA2 or BicA2 proteins from *Prochlorococcus* spp. have not yet been described. **(C)** Phylogenetic analysis of BicA proteins from cyanobacteria reveal two BicA families [BicA1 and BicA2; 26]. Each is a member of the larger SulP transporter family [27]. Members of the BicA1 clade from *Picosynechococcus* PCC7002 (*Picosyn* PCC7002) and *Synechocystis* PCC6803 (*Syn* PCC6803; both identified in the BicA1 clade as white circles with coloured outline) have been previously characterised to show HCO_3_- transport function [17], however no function has yet been described for members of the BicA2 clade. Reported in this study is a functional analysis of BicA2 proteins from *Prochlorococcus* CCMP1375 and *Prochlorococcus* CCMP1986 (identified in the BicA2 clade as white circles with coloured outline). Note that some species of cyanobacteria (e.g. *Cyanobium* PCC7001 [*Cya* PCC7001]) contain multiple genes for BicA1 and BicA2 proteins. Sequence alignment performed using MAFFT 7.526 [38] from a curated BicA dataset containing 223 protein sequences (Supplementary data) with default parameters and 1000 bootstrap replicates. Tree graphics were generated using iTol [40]. Tree scale is substitutions per amino acid.

Currently, cyanobacterial CCMs broadly have a total of six identified inorganic carbon (Ci; CO_2_ and HCO_3_^-^) transport systems (Figure 1), classified into two main types; those that transport HCO_3_^-^ from outside the cell [BicA1, SbtA1, SbtA2, BCT1; 15,16-19] as well as specialized CO_2_-to-HCO_3_^-^ conversion systems (NDH-I_3_ and NDH-I_4_) that are located on internal thylakoid membranes [20–23]. Cyanobacterial HCO_3_^-^transporters play a fundamental role in carbon assimilation in aquatic environments, since HCO_3_^-^ is the dominant form of Ci available in seawater and it has relatively slow diffusion across biological membranes compared with CO_2_ [24,25]. Both HCO_3_^-^ and CO_2_ uptake systems are found in most cyanobacterial species, except for *Prochlorococcus* spp. which do not possess genes for the known CO_2_-to-HCO_3_^-^conversion systems [26,27]. In addition, *Prochlorococcus* spp. generally only possess SbtA2 homologues and distant relatives of BicA1, which have been termed BicA2 [20,26,28]. SbtA2 was recently described as an apparent Cl^-^/SO_4_^2-^-dependent HCO_3_^-^transporter when SbtA2 proteins from *Synechococcus* spp. were heterologously expressed in *E. coli* [19]. However, the same study showed that SbtA2 homologues from *Prochlorococcus* spp. were inactive in this system, such that their physiological role remains uncertain. Despite the significance of this genus in global primary productivity [2,4] and evidence of CCM function [29,30], there has been no functional characterisation of Ci transport in *Prochlorococcus* spp. to date, leaving the mechanism of inorganic uptake in this important group unresolved.

Here we describe the functional characterisation of several BicA2 transporters from the globally dominant *Prochlorococcus* clade to confirm candidate proteins involved in HCO_3_ ^-^ uptake. *Prochlorococcus* spp. BicA2 genes were expressed in *E. coli* and screened for transport activity. BicA2 from the high-light ecotype *P. marinus* CCMP1986 showed no activity in its native form; however, adaptive laboratory evolution [15] produced a variant with a single point mutation that transported HCO_3_^-^in *E. coli*. Comparative sequence analysis of this mutated, functional form of BicA2 highlighted conserved amino acid sequence motifs near this mutation, guiding the identification of a BicA2 from the low-light ecotype *P. marinus* CCMP1375, which exhibited Na^+^-dependent HCO_3_^-^ transport in *E. coli*. These findings provide the first direct evidence of inorganic carbon transport by membrane proteins from *Prochlorococcus* spp., revealing a critical pathway for carbon entry into the biosphere.

## Materials and methods

### bicA2 gene cloning and expression construct assembly

BicA2 from *P. marinus* CCMP1986 (hereafter BicA2-1986) was amplified by PCR (Phusion High-Fidelity DNA Polymerase Thermo Fisher Scientific) from genomic DNA using primers listed in Table S1 and cloned (with and without mycH6 tag) using type IIS restriction enzyme Loop Assembly [31,32], under the control of the tetracycline inducible/repressible promoter and the *rrnB* terminator in the pCk1 (Kanamycin resistant) vector. BicA2 from *P. marinus* CCMP1375 (hereafter BicA2-1375) and its variants were synthesised by Synbio Technologies (USA) and cloned similarly to BicA2-1986, except the *lacI^q^-trc* promoter was used for expression experiments due to initial tests showing this was a more reliable promoter for this protein. Construct sequences were verified using Sanger sequencing (Macrogen Inc, Seoul, South Korea).

### HCO_3_^-^ uptake experiments using silicone oil centrifugation-filtration

HCO_3_^-^ uptake was examined in *E. coli* DH5α expressing putative transporters along with appropriate controls as per [33] with modifications. Starter cultures were grown overnight from glycerol stock (BicA2-1375 and variants) or from LB + 1.5% agar that had been growing for 72 h at 22°C (BicA2-1986 and variants). Expression was induced by the addition of isopropyl β-D-1-thiogalactopyranoside (IPTG) to a final concentration of 1 mM, or anhydrotetracycline hydrochloride (aTc; Merck) to 400 nM. Cells were depleted of Ci by washing three times with 1 mL of CO_2_-free buffer (20 mM BisTrisPropane-H_2_SO_4_ pH 7.5 + 200 mM NaCl + 10 mM K_2_HPO_4_). Due to the density of the silicone oil used, the maximum concentration of NaCl assessed in this assay was 200 mM. Assay buffer variations were used to test ion dependence of transporters, as specified.

### Complementation of CO_2_-dependent CAfree *E. coli*

*E. coli* CAfree strain [34] containing *bicA2* gene expression plasmids were grown overnight in LB + 20 mM HEPES-KOH pH 8.0 with antibiotics at 37°C in air supplemented with 4% CO_2_. These were diluted 1:5 in LB + 20 mM HEPES-KOH Ph 8.0 with antibiotics and grown for approximately 60 min. Culture densities (OD_600_) were measured and standardised before spot-plating onto Luria-Bertani (LB) agar containing ∼170 mM NaCl, 1.5% Bacto Agar and 20 mM HEPES-KOH pH 8.0, supplemented with kanamycin (50 µg.mL^-1^) and 400 µM isopropyl IPTG (for plasmids containing the *lacI^q^-trc* promoter) or 200 nM aTc [35]. Plates were incubated for 24 - 48 h at 37°C in air levels of CO_2_ (∼0.04%) or 4% CO_2_ v/v in air.

### Membrane enrichment and WB test for tagged proteins

Samples were taken from induced *bicA2* gene expression cultures of *E. coli* used for Ci uptake experiments, pelleted and stored frozen at −20°C until membrane enrichment was performed as follows. Frozen pellets were resuspended in 0.7 mL of lysis buffer (50 mM HEPES-KOH pH 8.0, 100 mM NaCl, 5% glycerol) supplemented with ∼30 kU rLysozyme (Merck) and incubated at room temperature with inversion for 30 min. Cells were then bead-beaten for 3 min with ∼200 µL glass beads at maximum speed (BioSpec mini-beadbeater-16). Cells debris removed by centrifugation (20,238 × *g*, 30 s). Supernatant was transferred to fresh tubes and centrifuged at 16,700 × *g* for 30 min at 4°C. The supernatants (soluble fraction) were retained and pellets were resuspended overnight in 50 µL lysis buffer including 1% DDM (Merck) at 4°C. Insoluble material in this membrane-enriched fraction was removed by centrifugation (10 min, 20,238 × *g* 4°C). DDT (50 mM) and SDS-PAGE loading buffer were added to both soluble and membrane-enriched fractions, heated for 5 min at 90°C, and loaded onto NuPAGE 4-12% Bis-Tris SDS-PAGE gels. Separated proteins were subsequently transferred to PVDF membrane (Merck-Millipore) using Transblot Turbo, (BioRad). After blocking with 5% skim milk powder in Tris buffered saline, membranes were probed with anti-myc antibody (cat # M4439, Merck) and AP-conjugated-secondary antibody (cat # A3562, Merck), and developed with Attophos (Promega Corporation). Blots were visualised using the Chemidoc MP imaging system [BioRad; 36].

### Adaptive laboratory evolution

In our hands CAfree *E. coli* is unable to grow at 4% CO_2_ in M9 minimal medium unless supplemented with 1% LB. A starter culture of CAfree carrying plasmid pCk1-BicA2-1986 was first grown from glycerol stock in LB containing 50 µg.mL^-1^ kanamycin at 37°C in air supplemented with 4% CO_2_ for approximately 18 h. This starter culture was then diluted 1:50 in 5 mL M9 medium containing 1% LB supplemented with 50 µg.mL^-1^ kanamycin and 50 nM aTc, and grown at 37°C in air supplemented with CO_2_ to 1%. Once the cultures grew dense overnight at these permissive CO_2_ conditions they were diluted 1:100 and incubated in air levels of CO_2_ (0.04%) and continued as batch dilution cultures until the culture grew dense overnight. Cells from cultures that grew in air levels of CO_2_ overnight (100 µL), were plated on to LB + 1.5% agar supplemented with kanamycin and aTc, and growth was assessed in air. Ten independent isolates from air-grown plates were cultured and pDNA was prepared using miniprep spin column (Qiagen) for sequencing (Macrogen Inc, Seoul, South Korea). To ensure the mutation conferring growth in air was not due to a mutation in the background strain of CAfree, purified plasmids were reintroduced to CAfree cells and assessed for growth in air.

### BicA multiple sequence alignment and phylogenetic tree construction

Using identified amino acid sequences; GAGATMGTV and GAGATMRTV as BLAST queries [37], we recovered a total of 233 BicA protein sequences from cyanobacteria with either of these motifs conserved. MSA was performed using MAFFT 7.526 [with default parameters and 1000 bootstrap replicates; 38] after curating the BicA sequence dataset [39] to remove incomplete or duplicate sequences.

Trees (Figure 1; Supplementary Figure S2) were visualised and edited using iTol [https://itol.embl.de/; 40]. Clade designation to either BicA1 or BicA2 was initially confirmed after visual inspection of the tree and sequence logos were subsequently generated using separate alignments of the BicA1 and BicA2 sequences using WebLogo3 [https://weblogo.threeplusone.com/create.cgi; 41].

## Results

Phylogenetic analysis demonstrated that there are multiple BicA subfamilies across cyanobacterial clades [Figure 1; 26,27]. Here, we refer to the previously characterised BicA proteins [17,18,42] as BicA1, and the other main family as BicA2 (Figure 1), consistent with the terminology introduced by Rae *et al* [26]. Notably, several species possess at least two BicA proteins, including *Cyanobium* PCC7001, which appears to have two BicA1 proteins and one BicA2 protein ranging between 43% and 65% sequence identity to each other (Figure 1; Supplementary Figure S1). Since *Prochlorococcus* possesses only BicA2 and SbtA2 proteins [Figure 1; 29], and SbtA2 from *Prochlorococcu*s were not functional in *E. coli* [19], we considered that BicA2 from *Prochlorococcus* should display HCO_3_^-^ transport function.

To assess HCO_3_^-^ transport by candidate BicA2 proteins, we exploited a mutant strain of *E. coli*, CAfree. CAfree lacks all cellular CA enzymes [14,15,19,33,34], and is unable to survive at atmospheric CO_2_ concentrations (∼0.04% v/v CO_2_ in air) due to its inability to rapidly interconvert CO_2_ and HCO_3_^-^. The latter is required for small molecule and fatty acid biosynthesis via HCO_3_^-^-dependent carboxylases [43] and the low HCO_3_^-^ supply in CAfree under ambient CO_2_ leads to growth inhibition [15,19]. However, CAfree can grow in elevated CO_2_ (4% v/v in air) where bulk diffusion of CO_2_, and its slow interconversion with HCO_3_^-^, leads to sufficient HCO_3_^-^ to support one-carbon metabolism and growth. More importantly CAfree can survive in air levels of CO_2_ if an active HCO_3_^-^ transporter is present in the plasma membrane [15,16,19,33,34]. The constraints of CAfree to survive in ambient CO_2_ can be exploited as a screening system to test for functional HCO_3_^-^/CO_2_ transporters and carbonic anhydrases [14–16].

Expression of BicA2-1986 from *P. marinus* CCMP1986 in CAfree did not rescue cell growth in air levels of CO_2_ (Figure 2A). To assess gain-of-function potential in this non-functional BicA2 form, we carried out adaptive lab evolution in CAfree cells under permissive CO_2_ supply [15]. Here, we have taken advantage of the naturally occurring mutation rate of *E. coli* [approximately 1 in 10^9^-10^10^ nucleotides per generation; 44] to evolve function in a non-functional HCO_3_^-^ transporter. This was achieved by growing CAfree expressing non-functional BicA2 proteins in growth-permissive CO_2_ levels (∼1% CO_2_ v/v in air). Under these conditions, populations that arose from cells with evolved transporter function then dominated the population under low CO_2_ due to a growth advantage [15,19]. Two independent attempts to evolve function in BicA2-1986 using adaptive laboratory evolution resulted in isolates carrying a point mutation (A>G at 925bp in the gene coding sequence) giving rise to an amino acid change, R309G. This functional form is hereafter referred to as BicA2-R309G-1986. Mutations in the CAfree background strain and on the plasmid backbone were excluded as causes for observed function by reintroducing this mutant *bicA2* into CAfree *E. coli*. This confirmed growth at air levels of CO_2_ was attributable to the mutated BicA2-R309G-1986 transporter.

**Figure 2:**
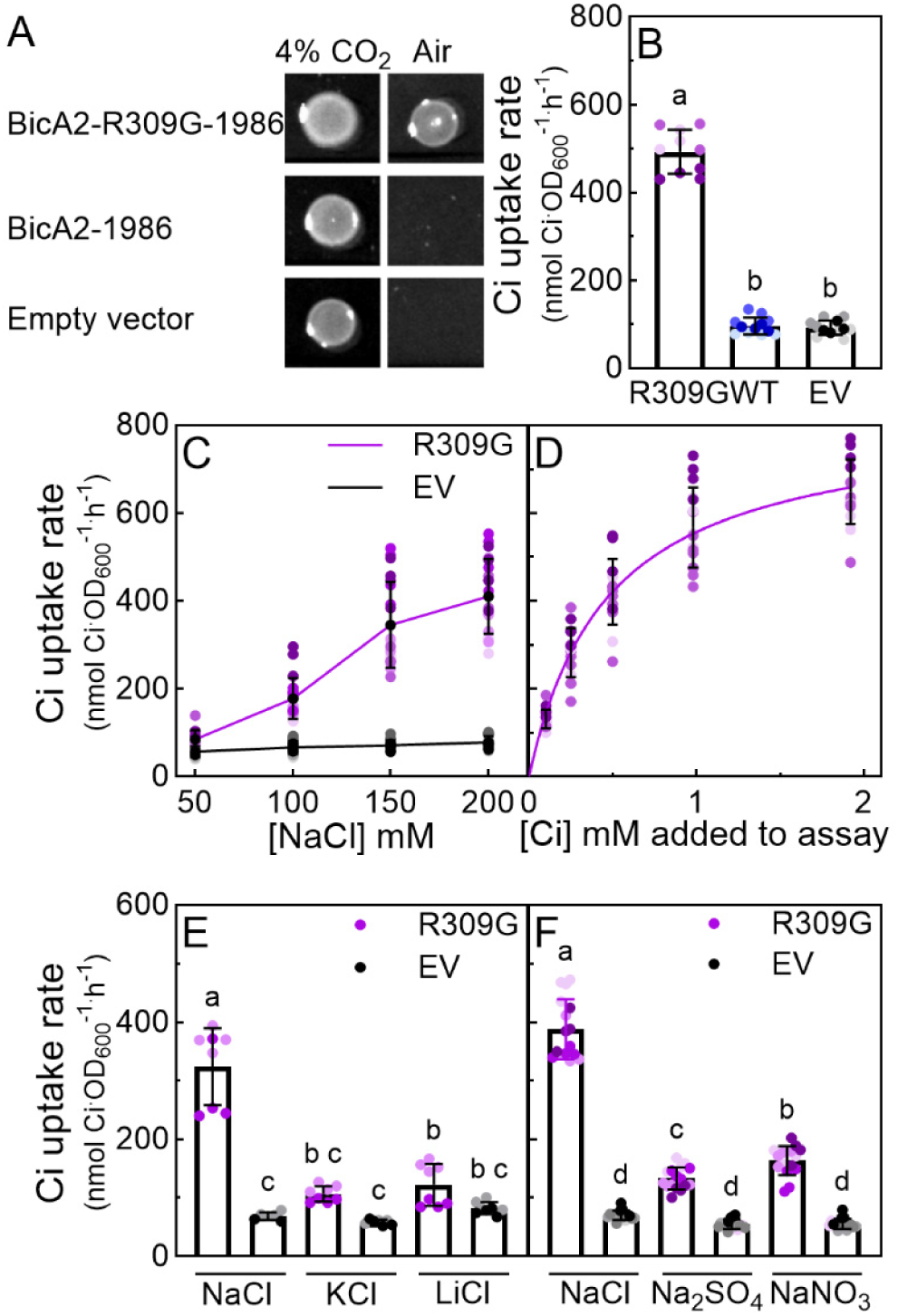
Characterisation of BicA2 forms from *Prochlorococcus* CCMP1986 via CAfree *E. coli* complementation and HCO_3_- uptake assays. **(A)** Cultures of CAfree *E. coli* expressing BicA2-R309G-1986, BicA2-1986 and a negative control (empty vector) were standardised for optical density (OD_600_) and spotted onto LB (pH 8.0) containing 1.5% agar supplemented with kanamycin 50 µg.mL_-1_ and 200 nM anhydrotetracycline. Plates were incubated at 37°C for 24 h in ambient (air) or 4% CO_2_. **(B)** The HCO_3_- uptake activity of *E. coli* DH5α expressing BicA2-R309G-1986 (R309G), BicA2-1986 (WT) and a negative control (empty vector; EV) were assessed. Cultures were inoculated from plates grown for 2 days at 22°C, and induced with 200 nM anhydrotetracycline for 2.5 h. Assay buffer composition was 20mM BTP-H_2_SO_4_ pH 7.5, 10 mM K_2_HPO_4_, and 200 mM NaCl. n=10-12 samples from three biological replicates. **(C)** The HCO_3_- uptake activity of *E. coli* DH5α expressing BicA2-R309G-1986 (R309G) and empty vector (EV) were assessed in response to increasing NaCl concentration. Assay buffer composition was 20 mM BTP-H_2_SO_4_ pH 7.5, 10 mM K_2_HPO_4_, (50, 100, 150 or 200 mM NaCl). Data are presented as mean ± s.d.; n=11 samples from three biological replicates. **(D)** The HCO_3_- uptake activity of *E. coli* DH5α expressing BicA2-R309G-1986 was assessed over a range of NaHCO_3_ concentrations (0.1-2 mM) to determine the affinity for HCO_3_-. For kinetic parameter estimates, the empty vector rates were subtracted prior to non-linear-regression to estimate the Michaelis-Menten constants (Table 1). Data are presented as mean ± s.d. with empty vector rates for each condition subtracted; n=14-16 samples from four biological replicates. Assay buffer composition was 20mM BTP-H_2_SO_4_ pH 7.5, 10 mM K_2_HPO_4_, and 200 mM NaCl. **(E)** The HCO_3_- uptake activity of *E. coli* DH5α expressing BicA2-R309G-1986 (R309G) and empty vector (EV) was assessed in the presence of NaCl, LiCl or KCl. Assay buffer composition was 20 mM BTP H_2_SO_4_ pH 7.5, 10 mM K_2_HPO_4_, 10 mM Na_2_HPO_4_, with relevant salts (150 mM NaCl or 150 mM KCl or 150 mM LiCl). Data are presented as means ± s.d.; n=6-8 samples from two biological replicates. **(F)** The HCO_3_-uptake activity of BicA2-R309G-1986 (R309G) and empty vector (EV) was assessed in response to different Na_+_ salts. Assay buffer composition: 20mM Hepes pH 7.5, 10 mM K_2_HPO_4_, and each of the salts (150 mM NaCl or 75 mM Na_2_SO_4_ or 150 mM NaNO_3_). Data are presented as means ± s.d. Different letters indicate significantly different means when assessed using one-way ANOVA (*p*<0.05).

**Table 1:** Kinetic parameters of BicA2 proteins heterologously expressed in *E. coli* DH5α.

| Transporter | $K_{0.5}$ (μM) | $V_{\max}$ (nmol Ci OD<sub>600</sub><sup>-1</sup> h<sup>-1</sup>) |
| --- | --- | --- |
| BicA2-R309G-1986 | 470 ± 42 | 812 ± 89 |
| BicA2-R309G-mycH6-1986 | 450 | 1690 |
| BicA2-1375 | 670 ± 180 | 320 ± 94 |
| BicA2-mycH6-1375 | 790 | 370 |

Data for untagged BicA2 forms is presented as mean ± s.d. of $K_{0.5}$ and $V_{\max}$ calculated from individual experiments where Michaelis-Menten non-linear fit had $r^2 > 0.8$ . Data for mycH6 tagged BicA2 are from single biological experiments.

To characterise BicA2-R309G-1986 function, HCO_3_^-^ uptake rates were determined in *E. coli* DH5α expressing both the mutant and WT transporter forms. This was assessed using H^14^CO_3_^-^ uptake assays employing the silicone oil centrifugation-filtration method [19]. Elevated HCO_3_^-^ uptake rates were observed in *E. coli* DH5α expressing BicA2-R309G-1986 when compared to those expressing either BicA2-1986 or an empty vector control, in the presence of 200 mM NaCl (Figure 2B). Typically, after subtraction of the empty vector control rates, HCO_3_^-^ uptake rates of approximately 500 nmol Ci.OD_600_^-1.^h^-1^ were observed in *E. coli* DH5α cells expressing BicA2-R309G-1986 (Figure 2B, C).

Since the predicted protein structure of the transmembrane portion of BicA2 shows a high degree of similarity to the transmembrane portion of the known Na^+^-dependent HCO_3_^-^ transporter BicA1 from *Synechocystis* PCC6803 (BicA1-6803^TM^; Figure 3), the HCO_3_^-^ uptake of *E. coli* DH5α expressing BicA2-R309G-1986 was assessed for Na^+^-dependence. BicA2-R309G-1986 HCO_3_^-^ uptake rates increased in response to elevated NaCl concentrations, up to the maximum concentration possible (200 mM NaCl) in the silicone-oil assay (Figure 2). In the presence of alternative cations, K^+^ and Li^+^, BicA2-R309G-1986 demonstrated similar HCO_3_^-^ uptake activity as the empty vector control (Figure 2), suggesting that Na^+^ was the effective cation for HCO_3_^-^ transport. However, when assessed in the presence of various Na^+^ salts where the Na^+^ concentration remained constant, the HCO_3_^-^ uptake activity of BicA2-R309G-1986 was variable (Figure 2), suggesting an alternative control mechanism to Na^+^ alone.

**Figure 3:**
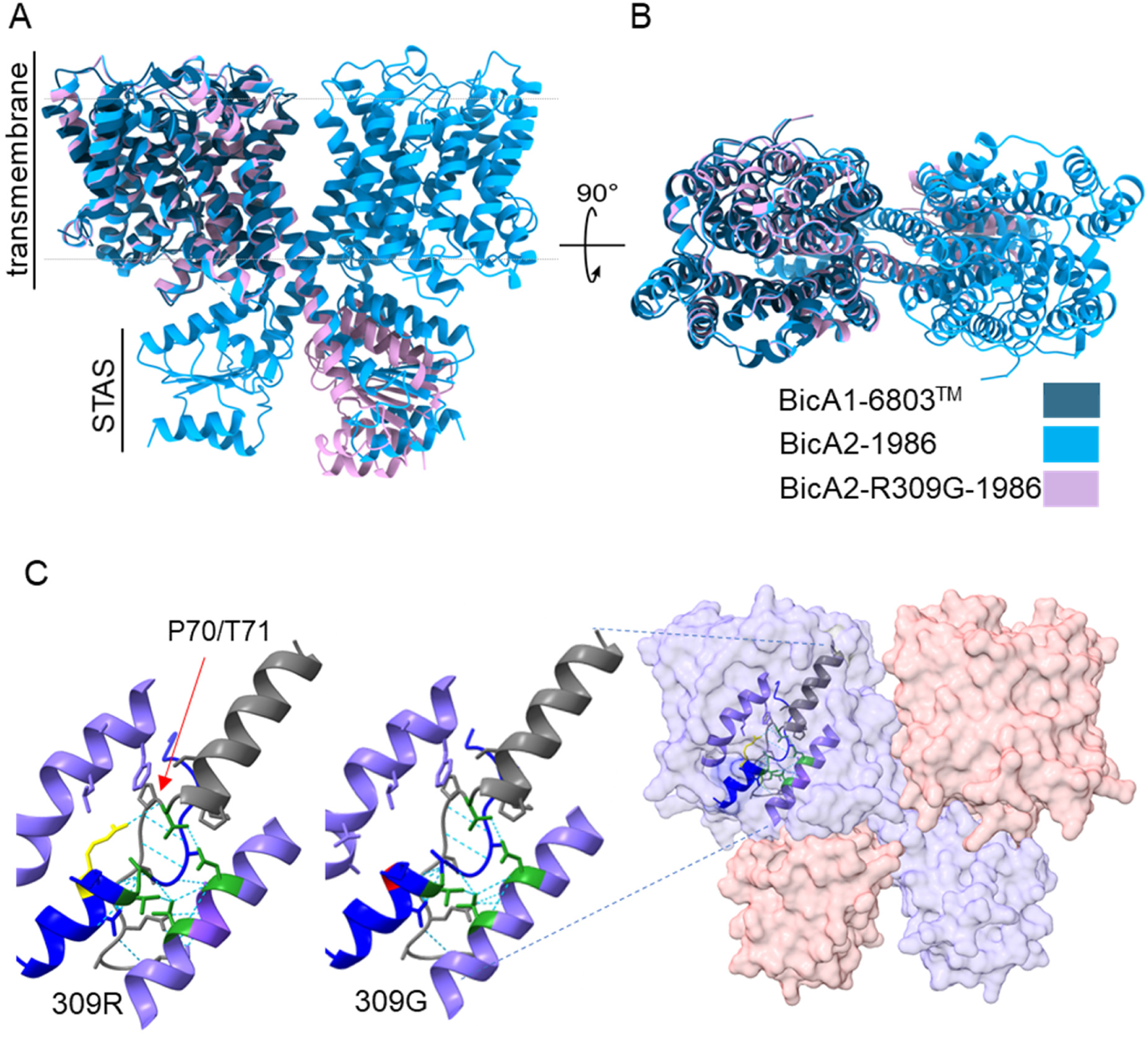
Comparison of predicted protein structures of BicA2-1986 variants and BicA1-6803^TM^. **(A)** The predicted strutures of BicA2-1986 homodimer and BicA2-R309G-1986 monomer [AlphaFold 3; 55] overlaid with the transmembrane portion of BicA1 from *Synechocystis* PCC6803 [PDB: 6KI1; BicA1-6803_TM_; 42]. A side view with dashed line approximating the membrane, and **(B)** an extracellular view. **(C)** The molecular structure of the putative HCO_3_-binding area (crossover of TM 3, grey; and TM 10, dark blue) of BicA2-1986 and BicA2-R309G-1986 with position 309 highlighted to illustrate the difference between arginine and glycine; G309 (red), R309 (yellow), GAGATM[G/R]TV motif (dark blue) and putatitive HCO_3_-binding amino acids [green; 42]. Dotted lines indicate putative H-bonds. The putative H-bonds between R309 and P70/T71 are not predicted when G309 is present within the predicted ion-binding domain [green; 42]. Structural alignment, H-bond prediction and visualisation using ChimeraX [58].

The HCO_3_^-^ uptake of *E. coli* DH5α carrying BicA2-R309G-1986 was then assessed in response to a range of NaH^14^CO_3_ concentrations to estimate its affinity for HCO_3_^-^. The concentration of HCO_3_^-^ providing half maximal uptake rate (K_0.5_) was estimated to be 470 ± 42 µM, and a maximum uptake rate (V_max_) estimated on a culture density basis was 812 ± 89 nmol Ci.OD_600_^-1.^h^-1^ (Figure 2).

Alignments of BicA2 with BicA1 protein sequences that have previously shown HCO_3_^-^ transport function [17,18,42] highlighted a highly conserved motif GAGATM[G/R]TV in transmembrane (TM) helix 10 (Figure 3A; Supplementary Figure S2) associated with the observed R309G mutation in BicA2-1986 (Figure 3). Both BicA2-R309G-1986 and BicA1 proteins that have demonstrated HCO_3_^-^ transport function to date, have a glycine in this corresponding position (Figure 3; Supplementary Figure S1). We therefore reasoned that the G309 is important for HCO_3_^-^ transport function, and we hypothesised that naturally active forms of BicA2 would carry a glycine residue at this corresponding position. Therefore, in order to identify potentially active native forms of BicA2 in *Prochlorococcus* spp., BLAST searches were performed to identify sequences with high homology to BicA2-1986 (>80% identity) that contained a glycine residue in the conserved TM10 GAGATM**<u>G</u>**TV motif. A single candidate protein (accession #WP_011124397.1) was identified from *P. marinus* CCMP1375 (BicA2-1375) that contained GAGTM**<u>G</u>**TV in TM10 and had high sequence identity (86.8%) to BicA2-1986 (Supplementary Figure S1; Supplementary Figure S2).

CAfree *E. coli* cells expressing BicA2-1375 grew at air levels of CO_2_ (Figure 4A). However, when compared to BicA2-R309G-1986 (which complemented CAfree in air within 24 h; Figure 2A), CAfree expressing BicA2-1375 grew slower in air with growth only evident after 48 h (Figure 4A). HCO_3_^-^ transport by BicA2-1375 was confirmed by HCO_3_^-^ uptake assays (Figure 4B). Like BicA2-R309G-1986, HCO_3_^-^uptake by BicA2-1375 was also found to be Na^+^-dependent (Figure 4C, D) and showed little response to alternative cations (K^+^, Li^+^; Figure 4D).

**Figure 4:**
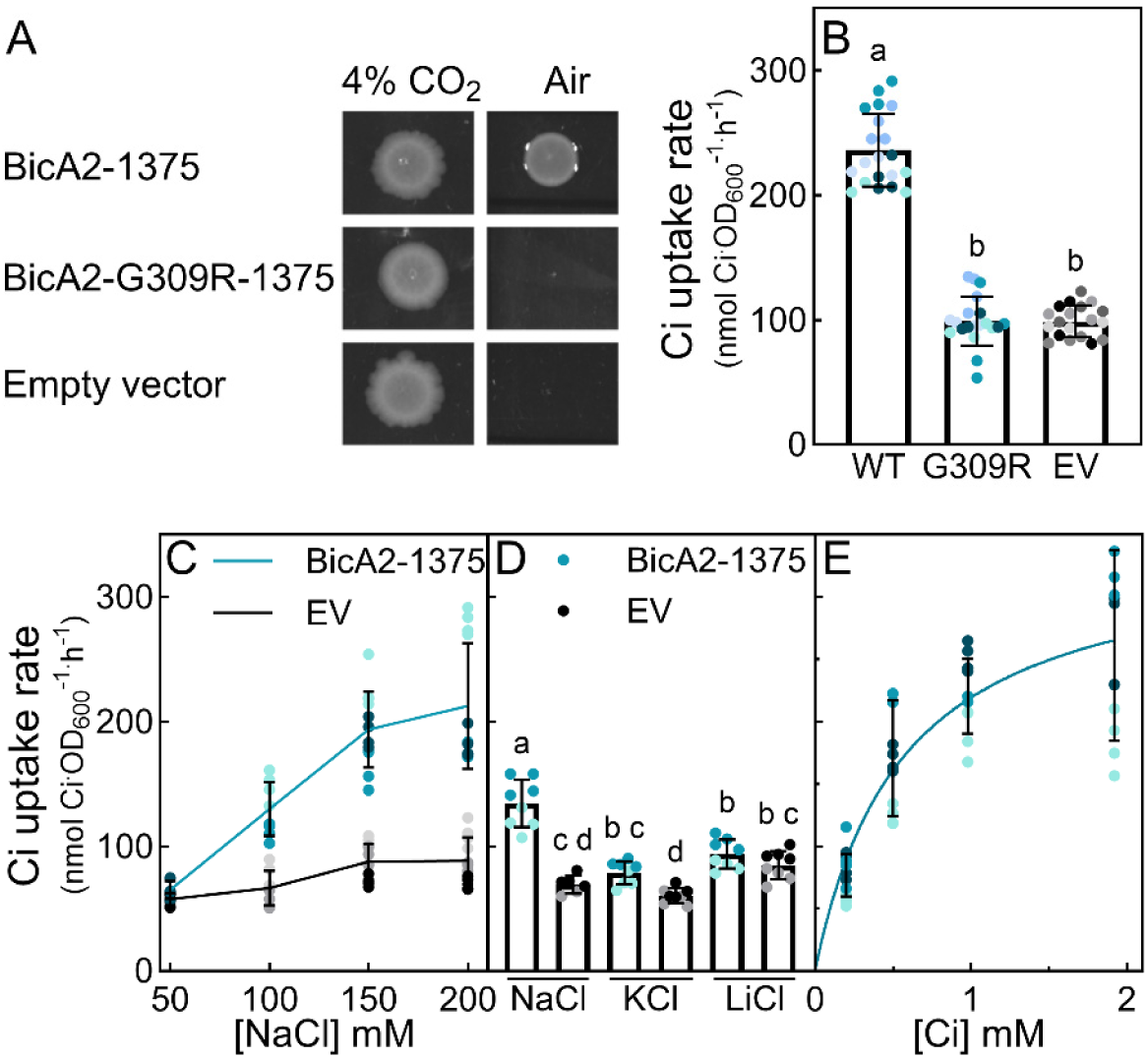
Characterisation of BicA2 forms from *Prochlorococcus* CCMP1375 via CAfree *E. coli* complementation and HCO_3_- uptake assays. **(A)** Cultures of CAfree *E. coli* expressing BicA2-1375, BicA2-G309R-1375 and a negative control (empty vector) were standardised for optical density (OD_600_) and spotted onto standard LB (pH 8.0) containing 1.5% agar supplemented with kanamycin 50 µg.mL_-1_ and 400 µM isopropylthiogalactopyranoside. Plates were incubated at 37°C for 48 h in ambient CO_2_ (air) and in 4% CO_2_ for 20 h. **(B)** The HCO_3_- uptake activity of *E. coli* DH5α expressing BicA2-1375 (WT), BicA2-G309R-1375 (G309R) and a negative control (empty vector; EV) was assessed. Cultures were induced with 1mM isopropylthiogalactopyranoside for 2.5 h. Assay buffer composition was 20 mM BTP H_2_SO_4_ pH 7.5, 10 mM K_2_HPO_4_, and 200 mM NaCl. Data are presented as means ± s.d. n=20 samples from five biological replicates. **(C)** The HCO_3_- uptake activity of *E. coli* DH5α expressing BicA2-1375 and empty vector (EV) was assessed in response to increasing NaCl concentration in the assay buffer. Assay buffer composition was 20 mM BTP H_2_SO_4_ pH 7.5, 10 mM K_2_HPO_4_, and NaCl (50, 100, 150 or 200 mM). Data are presented as mean ± s.d.; n=8-12 from two to three biological replicates. **(D)** The HCO_3_- uptake of *E. coli* DH5α expressing BicA2-1375 and empty vector (EV) was assessed in the presence of NaCl, LiCl or KCl. Assay buffer composition was 20 mM BTP H_2_SO_4_ pH 7.5, 10 mM K_2_HPO_4_, 10 mM Na_2_HPO_4_, with relevant salts (150 mM NaCl or 150 mM KCl or 150 mM LiCl). Data are presented as means ± s.d. n=8 from two biological replicates. Different letters indicate significantly different values when assessed using one-way ANOVA (*p*<0.05). **(E)** The HCO_3_- uptake activity of *E. coli* DH5α expressing BicA2-1375 was assessed over a range of NaHCO_3_ concentrations (0.25 - 2 mM) to determine the affinity for HCO_3_-. For kinetic parameter estimates, the empty vector rates were subtracted prior to non-linear-regression to estimate the Michaelis-Menten constants (Table 1). Data presented as mean ± s.d. with empty vector rates specific for each condition subtracted. n=12 from three biological replicates. Assay buffer composition was 20 mM BTP-H_2_SO_4_ pH 7.5, 10 mM K_2_HPO_4_, and 200 mM NaCl.

Silicone oil centrifugation-filtration assays were used to compare BicA2-1375 and BicA2-R309G-1986 HCO_3_^-^ uptake kinetics. BicA2-1375 showed similar affinity for HCO_3_^-^ to BicA2-R309G-1986, but lower HCO_3_^-^ transport flux (Figure 4B, E; Table1). Using estimated *E. coli* cell volumes from a previous study by our group [16], and measured HCO_3_^-^ fluxes (in the presence of 200 mM NaCl and 2 mM Ci), we estimated cellular Ci pool sizes achieved after 30 sec uptake assays were between 4.8 and 7.3 mM, and between 3.6 and 6.2 mM for BicA2-R309G-1986 and BicA2-1375, respectively (Table 2).

**Table 2:**
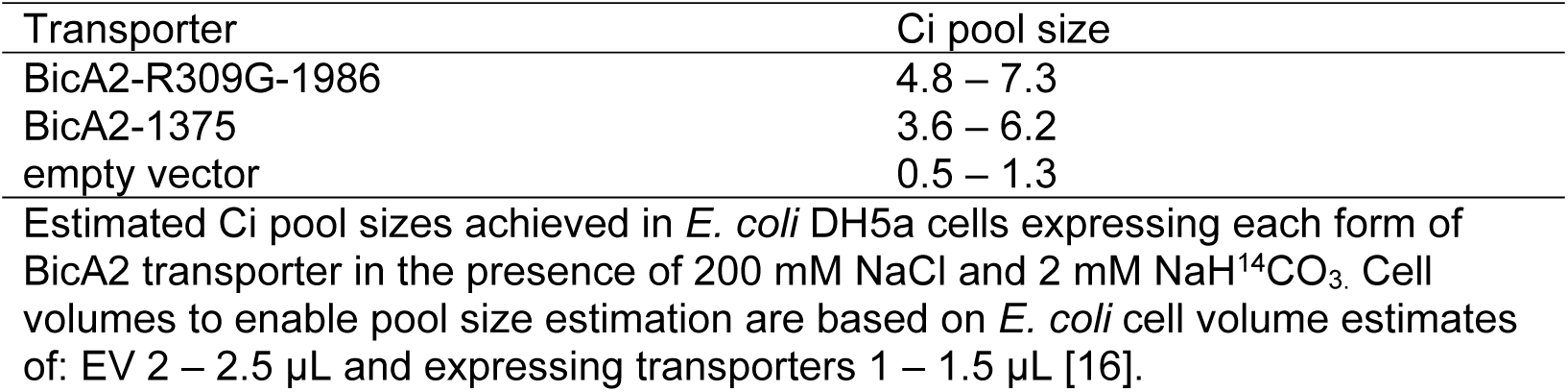
Estimated bicarbonate pool sizes in *E. coli* DH5α expressing BicA2 forms from *Prochlorococcus* spp.

We then tested the hypothesis that G309 (GAGATM**<u>G</u>**TV) was important to the activity of BicA2-1375 by generating a mutant form with arginine in this position (hereafter BicA2-G309R-1375), generating the same TM10 motif sequence as the non-functional WT form of BicA2-1986. This mutation abolished growth of CAfree *E. coli* at air levels of CO_2_ (Figure 4A), and BicA2-G309R-1375 showed no significant HCO_3_^-^ uptake above the empty vector control in *E. coli* DH5α (Figure 4B).

This outcome led us to explore BicA1 and BicA2 TM10 motif sequence conservation in greater detail through multiple sequence alignment (MSA). Using GAGATMGTV and GAGATMRTV as BLAST queries, we recovered a total of 233 BicA protein sequences with either of these motifs conserved. MSA and phylogenetic tree construction allowed us to identify that the GAGATM**<u>G</u>**TV motif is almost exclusively found in the BicA1 clade, while GAGATM**<u>R</u>**TV is almost exclusively found in the BicA2 clade (Supplementary Figure S2). This analysis also highlighted that, while BicA2-1986 and BicA2-1375 are closely related, BicA2-1375 was the only identified BicA2 sequence in our analysis that contained a BicA1-like GAGATM**<u>G</u>**TV motif (Supplementary Figure S2).

To identify the sub-cellular location of the BicA2 proteins when expressed in *E. coli* cells, and to compare the relative expression of mutant and WT BicA2 forms from *Prochlorococcus* CCMP1986 and CCMP1375, each protein was C-terminally tagged with a mycH6 epitope using a glycine and serine linker (GS-EQKLISEEDL-HHHHHH). The HCO_3_^-^ uptake activity of tagged forms of BicA2 was not impeded (Figure 5). Indeed, the tagged form of BicA2-R309G-1986 had approximately twice the HCO_3_^-^uptake rate as its untagged counterpart when measured based on cell density. This could be due to improved stability of BicA2-R309G-mycH6-1986 rather than a modulation of protein function (Figure 5A).

**Figure 5:**
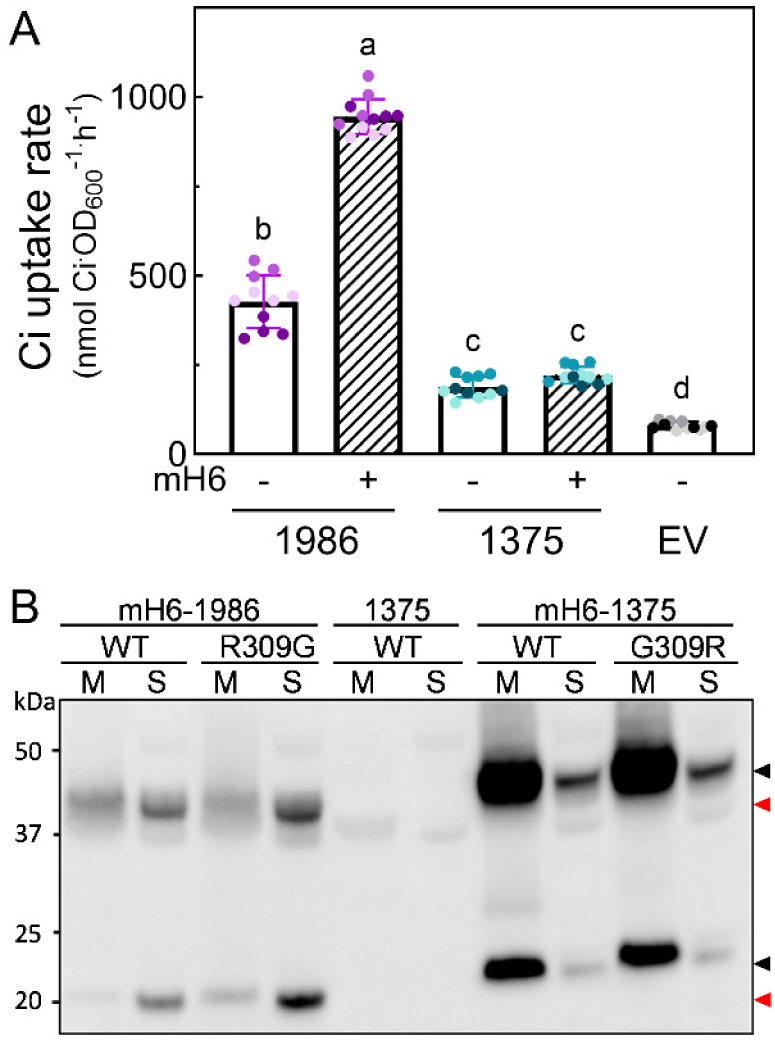
Comparison of protein abundance and HCO_3_- uptake activity of epitope (mycH6)-tagged BicA2 forms. **(A)** Comparison of HCO_3_- uptake activity of tagged (mH6) and untagged BicA2-R309G-1986 and BicA2-1375, and empty vector (EV) in *E. coli* DH5α. Assay buffer was 20 mM BTP-H_2_SO_4_ pH 7.5, 10 mM K_2_HPO_4_, and 200 mM NaCl. Data are presented as means ± s.d.; n=12. Different letters indicate significantly different values when assessed using one-way ANOVA (*p*<0.05). **(B)** Western blot analysis of crude soluble (S) and membrane-enriched (M) fractions of *E. coli* DH5α expressing BicA2-mycH6-1986 (WT-mH6-1986), BicA2-R309G-mycH6-1986 (R309G-mH6-1986), BicA2-mycH6-1375 (WT-mH6-1375), BicA2-G309R-mycH6-1375 (G309R-mH6-1375) and BicA2-1375 (WT-1375) to compare the abundance of mycH6-tagged proteins. Immunoblots were probed with anti-myc antibody. BicA2-1375 lacking a mycH6 tag is included as a negative control. Red arrowheads indicate BicA2-mycH6-1986 and black arrowheads indicate BicA2-mycH6-1375.

Using this epitope tag as a semi-quantitative tool, we assessed relative protein abundance by immunoblotting on soluble (cytoplasmic) and membrane-enriched *E. coli* DH5α cell fractions. We found that the majority of BicA2-R309G-mycH6-1986 was present in the soluble fraction (Figure 5B), suggesting minimal insertion and stability of this protein into the cell membrane. Furthermore, there was a large disparity between the relative quantities of BicA2-R309G-mycH6-1986 and BicA2-mycH6-1375 expressed in *E. coli* DH5α, indicating that, on a per-protein basis, BicA2-R309G-1986 is the more active form of the pair, or possibly less prone to protease breakdown at the C-terminus (Figure 5).

## Discussion

The dominant marine cyanobacterial genus *Prochlorococcus* [4] plays a critical role in global carbon fixation [2,45], yet its mechanism to take up Ci from the aquatic environment has, to date, been speculative. To our knowledge, this is the first report of functional HCO_3_ ^-^ uptake by a transporter originating from *Prochlorococcus*. *Prochlorococcus* spp. carry genes for distant relatives of the previously characterised HCO_3_ ^-^ transporters, SbtA1 and BicA1, namely SbtA2 and BicA2 [19,26,27]. However, the *Prochlorococcus* homologues represent distinct sub-families [Figure 1; 19,26,27] with potentially different modes of operation. While SbtA2 from *Synechococcus* spp. have shown HCO_3_^-^ transport activity [19], no verified bicarbonate uptake function has been reported for either SbtA2 or BicA2 proteins from *Prochlorococcus* spp.

Here we demonstrate HCO_3_ ^-^ transport by BicA2 homologues from *Prochlorococcus*, and identification of a conserved sequence motif in TM10 across BicA2 homologues. The active BicA2 proteins display low affinity, Na^+^-dependent HCO_3_ ^-^ transport (Figure 2; Figure 4; Table 1), and are the first to be identified in the ecologically important *Prochlorococcus* genus.

### HCO_3_^-^ transporters and their heterologous expression

The BicA class of transporters are part of the broader SLC26 sulfate transporter superfamily [17]. Proteins in this family typically consists of two domains, the N-terminal membrane transport domain and a regulatory sulfate transporter anti-sigma antagonist (STAS) domain constituting the C-terminal portion of the protein [42,46]. Like SbtA1 and SbtA2 [19], BicA1 and BicA2 appear to belong to distinct subfamilies [Figure 1; 26]. BicA1 has been characterised to be a Na^+^-dependent HCO_3_^-^ transporter with low substrate affinity, and a relatively high flux compared to SbtA1 and BCT1 [17,47]. Function of BicA1 has been described in its native host, *Picosynechococcus* PCC7002 [17], as well as when heterologously expressed in cyanobacteria [*Synechococcus* PCC7942; 17, *Synechocystis* PCC6803; 18], and the chemoautotrophic bacterium *Cupriavidus necator* H16 [48]. However, its functional characteristics have been elusive in *E. coli*, where function appears to be locked in an inactive state [16].

There has been no specifically observed function of any BicA2 forms to date, except for the physiological analysis of *Prochlorococcus* CCMP1986 [also known as *Prochlorococcus* MED4; 29] which has two proteins related to BicA1, including one closely related BicA2 form and a more distant relative [Figure 1; 27,29], and an SbtA2 [19,26]. After initial screening in *E. coli* CAfree showed no function, an evolved form of BicA2-1986 with a mutation in a conserved TM10 sequence motif, GAGATM**<u>G</u>**TV, allowed us to identify a sequence relative (BicA2-1375) that was functional in CAfree in its native form. This finding enabled identification of a conserved motif that plays a role in BicA2 function.

Functional characterisation of the two active BicA2 forms (BicA2-1375 and BicA2-R309G-1986) showed they both had K_0.5_ for HCO_3_^-^ of approximately 470-670 µM (Figure 2; Table 1). This is low affinity compared to estimates of K_0.5_ for HCO_3_^-^ for SbtA1 and SbtA2 of <100 - 190 µM, and ∼150 µM respectively when measured in *E. coli* [16,19]. Even though the affinity for HCO_3_^-^ of these BicA2 proteins is relatively low, these values still lie below the HCO_3_^-^ concentrations available in marine environments [∼2 mM HCO_3_^-^; 49]. This highlights the potential of BicA2 to function effectively at physiologically relevant conditions. In addition, estimated cellular Ci pool sizes of up to 6.2 mM and 7.3 mM (BicA2-1375 and BicA2-R309G-1986 respectively; Table 2) were achieved in *E. coli* when 2 mM Ci was supplied in uptake assays. Noting this is an artificial system, these pool sizes approach the level required to achieve saturation of CO_2_ fixation (>200 µM CO_2_) of the Form-IA Rubisco found in these organisms within their α-carboxysomes [50,51] through equilibration of the cellular HCO_3_^-^ pool with CO_2_ inside the carboxysome [7]. Notably, Scott *et al* [30] suggest *P*. *marinus* MIT9313 likely has at least a high affinity HCO_3_^-^ transporter (possibly SbtA2). Hopkinson *et al* [29] provide evidence that the relatively simple CCM of *Prochlorococcus* CCMP1986 (consisting of at least one BicA2 protein and an SbtA2 as the only Ci uptake systems; Figure 1) operates efficiently. Internal Ci pools sizes in *Prochlorococcus* CCMP1986 were measured to be approximately 15 mM in the presence of 2 mM external Ci [29]. The data presented here support the idea that BicA2 accounts for a substantial fraction of Ci uptake in *Prochlorococcus*, representing a major pathway for carbon influx and contributing significantly to global carbon fixation.

When the Na^+^-dependence of BicA2-R309G-1986 was assessed in *E. coli*, the highest HCO_3_^-^ uptake rates were evident in the presence of NaCl, but not when equimolar quantities of Na^+^ were supplied as Na_2_SO_4_ or NaNO_3_ (Figure 2F). This suggests a potentially undescribed mechanism of action, and may indicate a parallel Na^+^-driven Cl^-^/HCO_3_ ^-^ exchange mechanism, as seen in some HCO_3_^-^ transporters of the SLC4 (solute carrier 4) family [52]. Further assessment of this observation is required to confirm the mode of action of BicA2 members, which could potentially be evaluated in *Xenopus* oocytes [53,54].

### Key amino acid sequences for BicA2 activity

Structural studies performed by Wang *et al* [42] identified amino acid residues involved in putative HCO_3_^-^ binding sites in BicA1 from *Synechocystis* PCC6803. Alphafold3-generated structural alignments [55] reveal a high degree of shared tertiary elements between BicA1 and BicA2 (Figure 3). Additionally, BicA1 and BicA2 proteins that have shown HCO_3_^-^ transport function [this study; 17,18,42] demonstrate conservation of the proposed HCO_3_^-^ binding site amino acids and the GAGATM**<u>G</u>**TV motif in TM10 (Supplementary Figure S1). These identify potentially critical sequence elements across all BicA proteins that are essential for function. Wang *et al* [42] suggests that sequential A and T residues which we find in the GAG**<u>AT</u>**MGTV motif are important in HCO_3_^-^ and metal binding. The BicA1-6803 G_304_ residue (GAGATM**<u>G</u>**TV) is also likely involved in coordinating Na^+^ [42]. Additionally, the same study noted that some Cl^-^-coupled HCO_3_^-^ transporters (e.g. some SLC4 and SL26 family Cl^-^/HCO_3_^-^ exchangers) contain an arginine at the amino acid position corresponding to position 304 of BicA1-6803 (i.e. GAGATM**<u>R</u>**TV), while some Na^+^-coupled HCO_3_^-^ transporters (e.g. SLC4 family Na^+^/HCO_3_^-^ cotransporters and BicA1) contained a glycine at this position (i.e. GAGATM**<u>G</u>**TV), providing further evidence for different modes of action between BicA1 and BicA2.

The role of R_309_ in the GAGATM**<u>R</u>**TV motif of native BicA2-1986 remains unclear, despite this sequence being conserved across BicA2 proteins (Supplementary Figure S2). One possibility is that transporters with this motif require activation and are normally inactive. In contrast, the R309G mutation may produce an “always-active” form, bypassing such regulation. Structural predictions suggest that R_309_ forms H-bonds with P_70_/T_71_ in the WT protein, interactions absent from BicA2-R309G-1986 (Figure 3), potentially increasing conformational flexibility and enabling activity without activation partners.

We propose that canonical BicA2 proteins bearing the GAGATM**<u>R</u>**TV motif are intrinsically capable of HCO_3_^-^ transport but are normally constrained by regulatory or activation requirements that are not met in *E. coli*. Under this model, adaptive laboratory evolution or rare natural substitutions that replace the TM10 arginine with glycine may generate an “always-active” transporter, thereby bypassing these constraints. We note that BicA2-1375 appears to be an outlier in the BicA2 clade, having a GAGATM**<u>G</u>**TV motif in TM10 when all its close relatives have a GAGATM**<u>R</u>**TV motif in this region (Supplementary Figure S2), yet it is functional while BicA2-1986 (having the canonical motif sequence for this clade) is not (Figure 4). It is possible that BicA2-1375 has evolved toward a canonical BicA1-type protein. It is conceivable that stable growth conditions for *Prochlorococcus* CCMP1375 in lab culture prior to its genomic sequencing [5], could lead to stable populations with this genotype.

We were able to generate a double mutant of the BicA2 protein from *Cyanobium* PCC7001 which gained function in the CAfree system, however, this form displayed very low rates of uptake (Supplementary Figure S3). Notably, this protein is from a cellular system complicated by the presence of; SbtA1 [16], a functional BicA1 form [17; Figure 1], and an additional BicA1 with no GAGATM motif (Figure 1, Supplementary Figure S2; Rae et al., 2011; Cabello-Yeves et al.) such that its functional role may be more complex than that described here for *Prochlorococcus*.

### Conclusion

*Prochlorococcus* spp. are major contributors to marine primary productivity [2], yet direct evidence for the HCO_3_^-^ transport pathways underpinning their photosynthesis has been lacking. Here, we demonstrate that BicA2 functions as a Na^+^-dependent HCO_3_^-^ transporter in *Prochlorococcus*. Together with the absence of alternative transport systems and its similarity to BicA1, our results support the conclusion that BicA2 is a functional HCO_3_^-^ transporter *in vivo*, likely providing a primary route of Ci uptake to sustain carboxysomal CO_2_ supply [29]. Our findings further suggest that BicA2 proteins are intrinsically capable of transport, with the conserved arginine in the TM10 GAGATMRTV motif imposing an inactive or regulated state that can be relieved by substitution to glycine. We propose that BicA2 operates alongside the higher-affinity SbtA2 [19] to enable efficient CCM function in this important contributor to global primary productivity.

## Acknowledgements

The authors thank David Savage (UC Berkeley) for kindly providing the CAfree strain of *E. coli*.

## Author contributions

LMR, GDP, BML: conceptualization and visualisation; GDP project administration; LMR, BML: methodology and analysis; BML, GDP, CSB: supervision; LMR: writing – original draft; all authors: writing – review and editing;

## Conflict of interest

The authors declare no conflicts of interest.

## Funding Statement

LMR received an Australian Government Research Training Program Scholarship to cover tuition fees.

## Data availability

The primary data supporting this study are publicly available at https://data.mendeley.com/datasets/g2f8ghkrnj/1 and are available from the corresponding author upon request. Additional data reported in this paper are presented in the Supplementary Data.

## Supplementary Information

Supplementary Table S1 Primers used for amplification of *bicA2-1986*

Supplementary Figure S1 Comparison of select BicA1 and BicA2 protein sequences demonstrate key sequence differences.

Supplementary Figure S2 GAGATM[G/R]TV motif conservation in transmembrane domain 10 (TM10) of BicA1 and BicA2 clades.

## Abbreviations

aTc: anhydrotetracycline hydrochloride
BicA: bicarbonate transporter A
CA: carbonic anhydrase
CAfree: specialized *E. coli* strain that lacks Cas
CCM: CO_2_-concentrating mechanism
CO_2_: carbon dioxide
Ci: inorganic carbon (CO_2_ and HCO_3_^-^)
HCO_3_^-^: bicarbonate ion
IPTG: isopropyl β-D-1-thiogalactopyranoside
LB: Luria-Bertani
Na^+^: sodium ion
WT: wild-type

## Notes

https://data.mendeley.com/datasets/g2f8ghkrnj/1

### Competing Interest Statement

The authors have declared no competing interest.

