## Supplementary data for "Heterologous expression in *E. coli* reveals the bicarbonate transporter BicA2 drives carbon uptake in marine *Prochlorococcus* spp"


| **Authors**: | **ORCID** | **ResearcherID** |
|---|---|---|
| Loraine M. Rourke [1,2] | 0000-0002-2600-8073 | OGM-9247-2025 |
| Caitlin S. Byrt [1] | 0000-0001-8549-2873 | I-9785-2018 |
| G. Dean Price [1] | 0000-0001-5906-4912 | C-9505-2009 |
| Benedict M. Long [1,3,4*] | 0000-0002-4616-2967 | B-3799-2008 |

**Addresses**:

[1]Plant Science Division, Research School of Biology, 134 Linnaeus Way, Australian National University, Acton, ACT, Australia, 2601 Current address: [2]Plant SynBio Australia, Plant Science Division, Research School of Biology, 134 Linnaeus Way, Australian National University, Acton, ACT, Australia, 2601,[3]Discipline of Biological Sciences, and [4]ARC Centre of Excellence in Synthetic Biology, School of Science, University Drive, The University of Newcastle, Callaghan, NSW, Australia, 2308


**This supplementary information includes:**

7 pages

1 table

2 figures

**Supplementary Table:**

| **Table S1 Primers used for amplification of *bicA2-1986*** | |
|---|---|
| 1986_BicA2R_pt1 | AAGAAGACCGGACACCACAGAATATGGAACAAGCGTT |
| 1986_BicA2F_pt1 | AAGAAGACAACTCAAGGTTTGGAAATAATTAATGGATTTAATC |
| 1986_BicA2R_pt2 | AAGAAGACAACTCGCGAACCGGCTAATTCATTTAAAGCAGCTTC |
| 1986_BicA2F_pt2 | AAGAAGACGGTGTCCGGATTTATGTCTGGCATAGGAG |

Each primer pair has been designed to enable domestication of the *Prochlorococcus marinus* CCMP1986 BicA2 sequence for Loop assembly [1], a silent change (GTC to GTG) removing a BsaI site within the coding sequence. Primers are designed 5' to 3'.

### Supplementary Figures

A

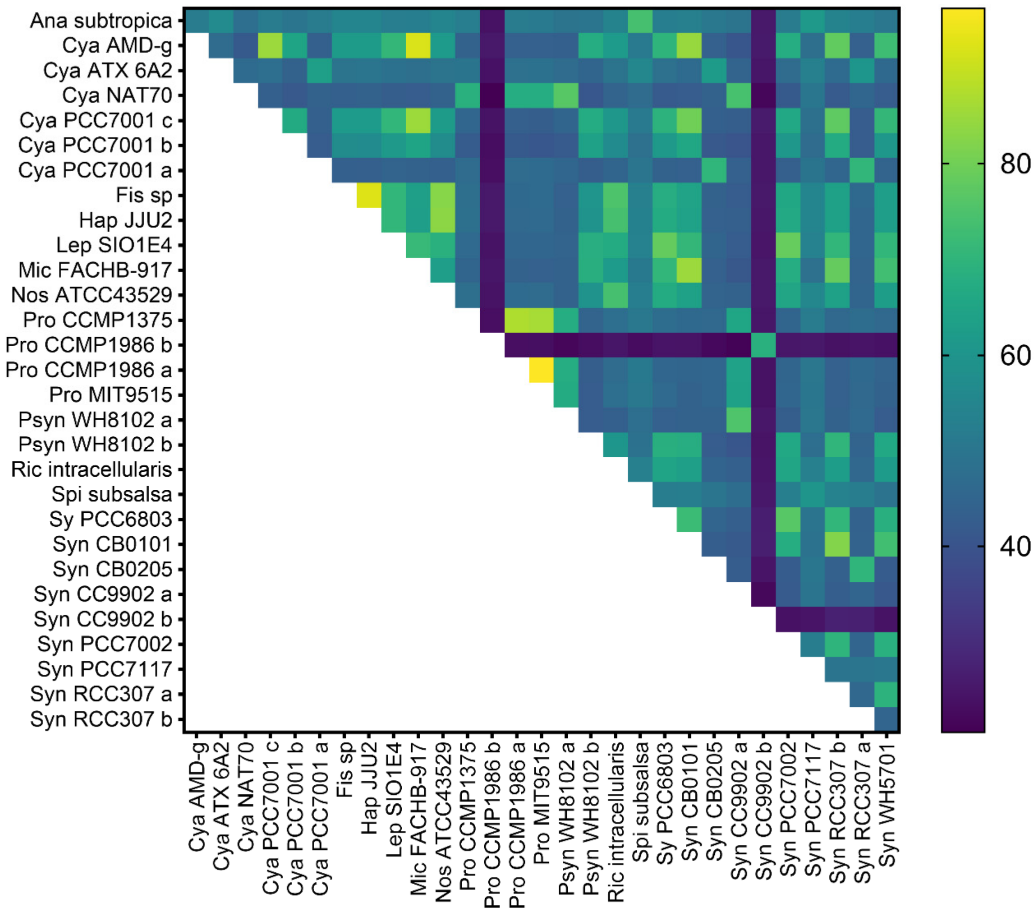

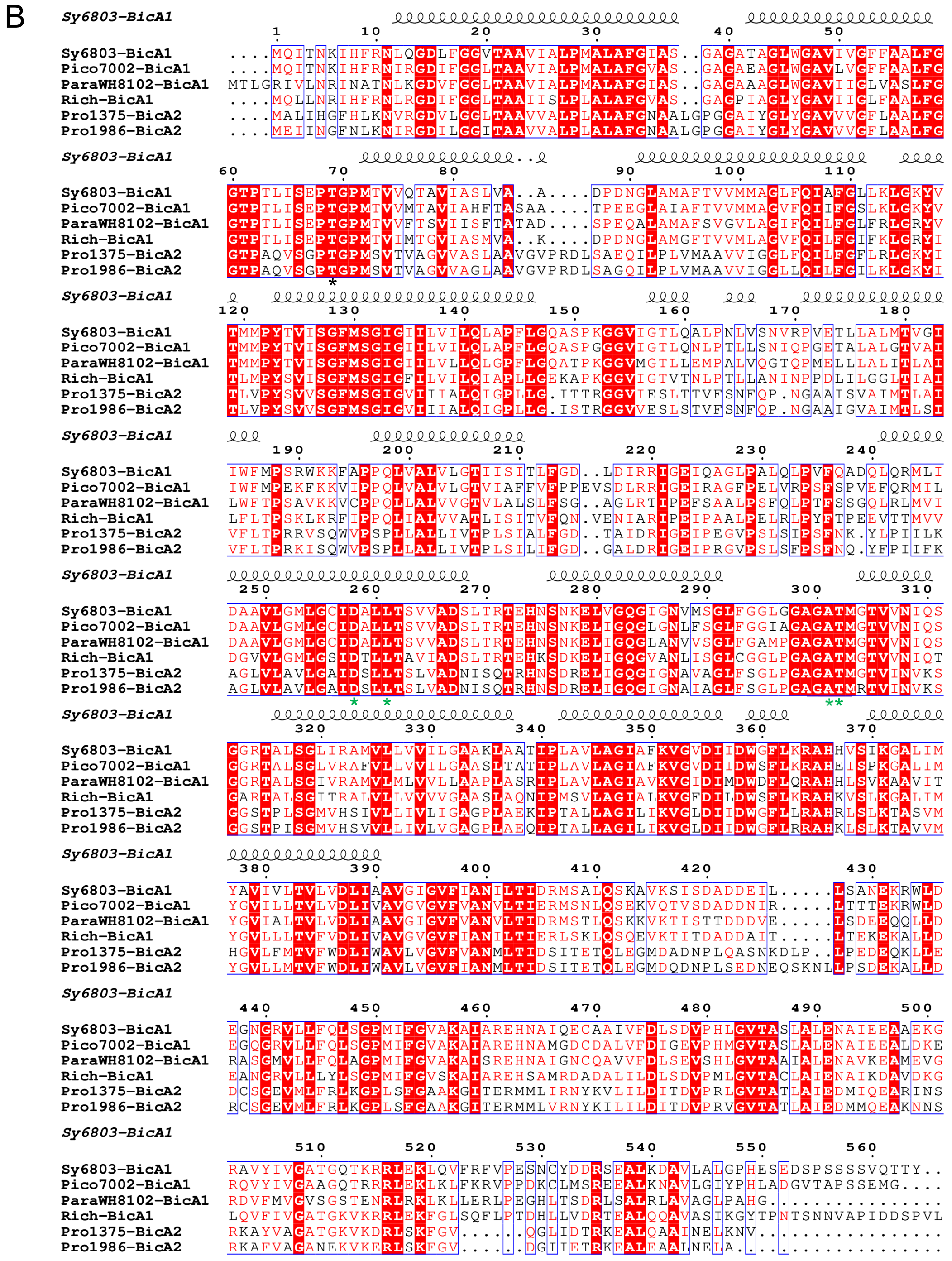


**Supplementary Figure S1: Comparison of select BicA1 and BicA2 protein sequences demonstrate key sequence differences.**

**A)** Heat map showing the % similarity of select members of the BicA family in Figure 1. The higher the number the more similar the sequence. Psyn: *Parasynechococcus*, Syn: *Synechococcus*, Pro: *Prochlorococcus*, Cya: *Cyanobium*, Mic: *Microcystis*, Lep:

*Leptolyngbya*, Fis: *Fischerella*, Hap: *Hapalosiphonaceae*, Nos: *Nostoc*, Sy: *Synechocystis*, Ana: *Anabaena*, Spi: *Spirulina*, Ric: *Richelia*, Pico: *Picosynechococcus.* Alignment performed by Geneious Prime and graphed using GraphPad. **B)** Alignment of the amino acid sequences of select members of the BicA family. See Figure 1 for the phylogeny tree. The structure for the transmembrane portion of *Synechocystis* PCC6803 BicA1 [2], was used for the structural label, alignment performed by MultAlin [3] and graphics by ESPript [4]. Asterisks (*) indicate the putative $HCO_3^-$ binding residues. The GAGATM[G/R]TV motif of transmembrane domain 10 (TM10) identified in this study begins at position 298 relative to the *Synechocystis* PCC6803 BicA1 sequence.

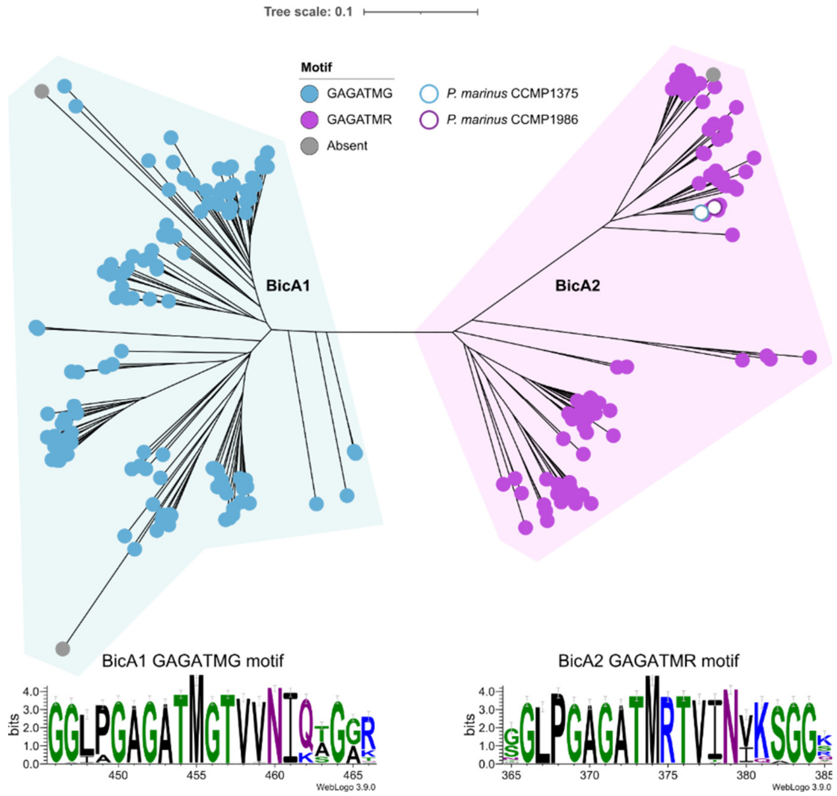


**Supplementary Figure S2: GAGATM[G/R]TV motif conservation in transmembrane domain 10 (TM10) of BicA1 and BicA2 clades.**

Identification of conserved GAGATMGTV or GAGATMRTV motifs across BicA forms highlights that the GAGTMGTV motif is almost exclusively found in the BicA1 clade, while GAGTMRTV is almost exclusively found in the BicA2 clade. The non-functional *Prochlorococcus marinus* CCMP1986 BicA2 form (BicA2-1986) has a GAGATMRTV motif in TM10 and is among relatives which also contain the GAGATMRTV motif. It's mutation to a BicA1-like GAGATMGTV sequence (BicA2-R309G-1986; Figure 2) enabled function. The natively functional BicA2 from *P. marinus* CCMP1375 (BicA2-1375) contains a GAGATMGTV motif in TM10 and its mutation to a BicA2-like GAGATMRTV motif (BicA2-G309-R-1375; Figure 4) leads to loss of function. Sequence alignment performed using MAFFT 7.526 [5] from a curated BicA dataset containing 223 protein sequences (Supplementary files) with default parameters and 1000 bootstrap replicates. Tree graphics were generated using iTol [6] identifying all sequences with a GAGATMGTV motif in blue, all GAGATMRTV motifs in purple and sequences with no identified GAGAMT[G/R]TV motif in grey. Tree scale is substitutions per amino acid. Sequence logos are based on separate alignments of the BicA1 and BicA2 sequences in the curated sequence dataset. Residues coloured by chemistry, and logos were generated using WebLogo3.

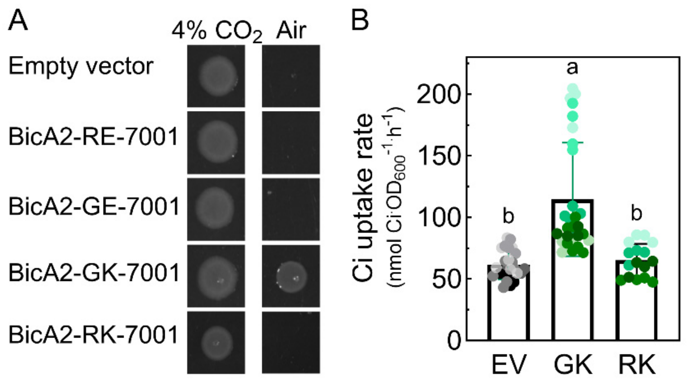


**Supplementary Figure S3**: **BicA2-7001 WT and mutants were assessed for $HCO_3^-$ transport using complementation and $HCO_3^-$ uptake assays**. Wild-type and mutant forms of BicA2-7001 containing GAGATM**R**$_{319}$TV and GAGAT**M**$_{319}$GTV respectively were assessed for $HCO_3^-$ transport in CAfree. Adaptive laboratory evolution of BicA2-R319G-7001 generated a form with functional $HCO_3^-$ transport activity BicA2-R319G,E370K-7001. **(A)** Complementation assays of CAfree assessed the function of BicA2-7001 forms with combinations of arginine or glycine at position 319, and glutamic acid or lysine at amino acid position 370. Cultures of CAfree carrying BicA2-7001 (BicA2-RE-7001), BicA2-R319G-7001 (BicA2-GE-7001), BicA2-R319G,E370K-7001 (BicA2-GK 7001), BicA2-E370K (BicA2-RK-7001) and pCK1 (empty vector) were standardised for optical density ($OD_{600}$) and spotted onto LB-agar pH 8.0 supplemented with kanamycin 50 µg.mL$^{-1}$ and 400 nM aTc. Plates were incubated at 37°C in ambient (~0.04% CO2) and 4% $CO_2$ for approx. 24 hours. **(B)** $HCO_3^-$ uptake activity of CAfree carrying BicA2-GK-7001 (GK) and BicA2-RK-7001 (RK) grown in air for the induction period and empty vector (EV) grown in 4% $CO_2$ during induction period. Assay buffer contained 20 mM BTP-$H_2SO_4$ pH 7.5 + 10 mM $K_2HPO_4$ + 200 mM NaCl. Data presented as mean ± s.d.; n=16-28 from 4-8 biological replicates. Data presented as means ± s.d. Different letters indicate significantly different means ($P< 0.05$) when analysed by one-way ANOVA.
